# Solving the influence maximization problem reveals regulatory organization of the yeast cell cycle

**DOI:** 10.1101/075069

**Authors:** David L Gibbs, Ilya Shmulevich

**Author notes:** Corresponding author: David L Gibbs^1^.

## Abstract

The Influence Maximization Problem (IMP) aims to discover the set of nodes with the greatest influence on network dynamics. The problem has previously been applied in epidemiology and social network analysis. Here, we demonstrate the application to cell cycle regulatory network analysis of Saccharomyces cerevisiae.

Fundamentally, gene regulation is linked to the flow of information. Therefore, our implementation of the IMP was framed as an information theoretic problem on a diffusion network. Utilizing all regulatory edges from YeastMine, gene expression dynamics were encoded as edge weights using a variant of time lagged transfer entropy, a method for quantifying information transfer between variables. Influence, for a particular number of sources, was measured using a diffusion model based on Markov chains with absorbing states. By maximizing over different numbers of sources, an influence ranking on genes was produced.

The influence ranking was compared to other metrics of network centrality. Although ‘top genes’ from each centrality ranking contained well-known cell cycle regulators, there was little agreement and no clear winner. However, it was found that influential genes tend to directly regulate or sit upstream of genes ranked by other centrality measures. This is quantified by computing node reachability between gene sets; on average, 59% of central genes can be reached when starting from the influential set, compared to 7% of influential genes when starting at another centrality measure.

The influential nodes act as critical sources of information flow, potentially having a large impact on the state of the network. Biological events that affect influential nodes and thereby affect information flow could have a strong effect on network dynamics, potentially leading to disease.

Code and example data can be found at: https://github.com/Gibbsdavidl/miergolf

**Author Summary:** The Influence Maximization Problem (IMP) is general and is applied in fields such as epidemiology, social network analysis, and as shown here, biological network analysis. The aim is to discover the set of regulatory genes with the greatest influence in the network dynamics. As gene regulation, fundamentally, is about the flow of information, the IMP was framed as an information theoretic problem. Dynamics were encoded as edge weights using time lagged transfer entropy, a quantity that defines information transfer across variables. The information flow was accomplished using a diffusion model based on Markov chains with absorbing states. Ant optimization was applied to solve the subset selection problem, recovering the most influential nodes.The influential nodes act as critical sources of information flow, potentially affecting the network state. Biological events that impact the influential nodes and thereby affecting normal information flow, could have a strong effect on the network, potentially leading to disease.

## Introduction

Living systems dynamically process information. Cell surface receptors capture information from the environment and relay the information by internal signaling pathways [1,2]. Information is transferred, stored, and processed in the cell via molecular mechanisms, often triggering a response in the regulatory program. These types of dynamic genetic regulatory processes can be modeled with network information flow analysis.

Network flows embody a general class of problems where some quantity flows from source nodes, across the edges of a graph, draining in sink nodes. Various forms of network flow methodologies have found success in algorithms such as Hotnet, ResponseNet, resistor networks, and others [3,4,5].

Recently, the influence maximization problem (IMP) has received a great deal of interest in social network analysis and epidemiology as a general method for determining the relative importance of nodes in a dynamic process [6,7]. The IMP aims to discover a set of source nodes that, after applying a diffusion model, covers as much of the network as possible [8,9]. Examples are found in modeling the spread of infectious disease in social networks and in identifying optimal targets for vaccination [10]. Propagation of infection does not follow algorithmically defined paths on graphs, i.e. shortest paths, but instead flows on all possible paths. Similar to quantities of virus, information can also be treated as a quantity flowing on networks [11,12,13].

A variant of ant optimization was used to solve the IMP. Although, ant optimization is best known in path optimization, it can also be applied to subset selection problems [14,15,16]. In ant optimization, ants construct potential solutions, as sets, which are scored and reinforced, encouraging good solutions in later iterations. In this work, the result of the optimization procedure is an optimal, or nearly optimal, set of nodes that maximizes network cover when applying the diffusion algorithm developed by Stojmirović and Yu [12]. In application to biological networks, the IMP essentially remains an unexplored area of research [17]. We then used the IMP to study the regulatory structure underlying the yeast cell cycle.

To understand topologically where the influential genes are situated, we compare the IMP solution sets to gene sets derived from other centrality metrics, such as degree centrality [18], betweenness-centrality where shortest paths are considered [19], and PageRank, where only incoming flows are considered [20].

The cell cycle process in Saccharomyces cerevisiae is well studied, but is not completely characterized [21]. Regardless, it is apparent that cell cycle regulation is controlled by a network that dynamically processes signals. From the bench, we have limited ways of observing the process, such as using gene expression data and cataloging the patterns of periodicity. To gain further understanding of the regulatory structure, we performed used time series data and publicly available regulatory databases to solve the IMP (Fig 1) [22,23].

**Figure 1.).**
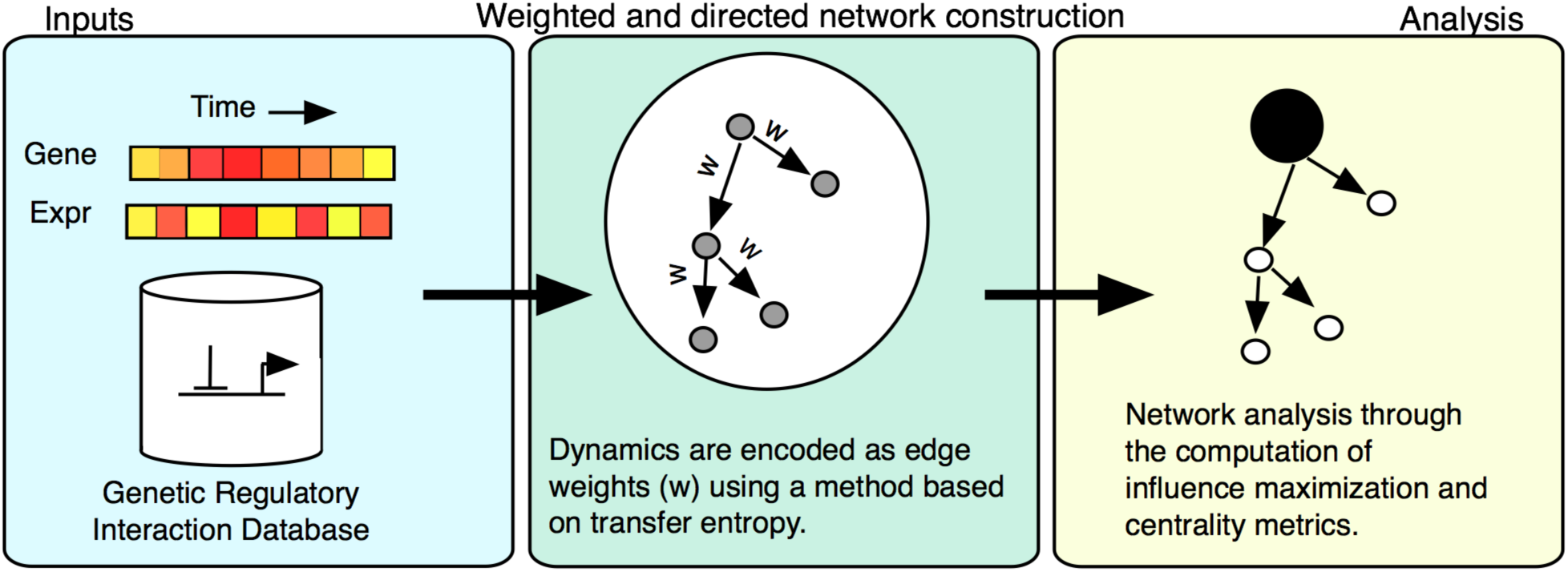
Analysis workflow. All regulatory edges from the YeastMine DB formed the regulatory network scaffold. Using time series gene expression data, time lagged transfer entropy was calculated and each edge was tested using a permutation-testing framework. The resulting network was used for solving the Influence Maximization Problem.

## Results

### Filtering regulatory edges using time lagged transfer entropy

Statistical metrics such as Pearson correlation are sometimes used to estimate the activity of regulatory edges. However, processes in biology do not instantaneously complete, and so various time lags are introduced to account for propagation time (SI Fig 1) [24]. Additionally, genetic regulatory interactions are directional; transcription factors act on genes, and not the other way around. So, although Pearson correlation is simple, there are more appropriate metrics to use with time series data, such as transfer entropy. Transfer entropy (TE) is a model-free method that attempts to quantify information transfer between two variables in a directional manner. Permutation-based statistics can be applied to assess the significance of TE. TE was computed over multiple time lags and summed. This avoids choosing a single time lag and is termed ‘sum-lagged transfer entropy’ or SLTE [35].

Using time series data for 5,080 measured genes and 26,827 genetic regulatory edges from YeastMine, both time-lagged Pearson correlation and sum-lagged transfer entropy was computed for all regulatory edges. Edges were removed if empirical p-values were greater than 1/*p*_*n*_ where *p*_*n*_ is the number of permutations (*p*_*n*_ = 50,000).

Time lagged Pearson correlation, where the maximum correlation is returned after considering a range of time lags (0-6 time steps), resulted in 7,729 edges, containing 3,216 nodes. Significant edge weights had a median correlation of 0.67. Most of the edges (76%) showed a maximum correlation when using a time-lag of zero.

The metric of interest, SLTE, resulted in 1,987 significant edges containing 1,147 nodes with median weights of 1.37 (Fig 2).

**Figure 2.).**
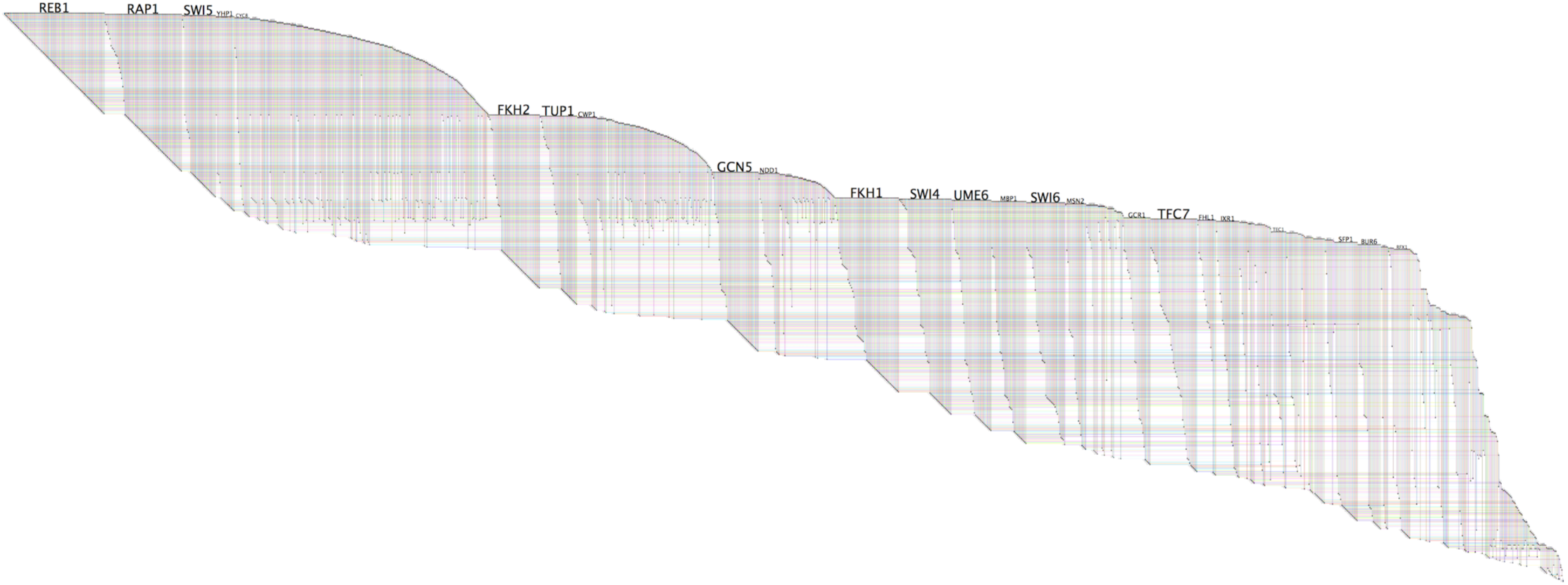
The resulting BioFabric network after significance testing. Nodes are shown as horizontal lines, with edges shown as vertical lines connecting nodes. High degree nodes can be seen as ‘wedges’ in the graph.

The overlap between the Pearson correlation and SLTE networks is moderate; only 12.8% of the edges in the correlation network are shared with the SLTE network (986 of 1,987 edges in the SLTE network or 49.6%), and while all SLTE nodes are found in the correlation network, only 33.9% of the correlation nodes are found in the SLTE network. When comparing the two different edge weights, Pearson’s and SLTE, on matched edges, the Pearson’s correlation between edge weights was low (0.39). Additionally, the mean node degree distribution in the correlation network is much higher than that of the SLTE network. For example, the SFP1 gene has degree 589 in the correlation network, compared to 60 in the SLTE network, summing both in-and out-edges. The high node degree in the correlation network suggests that correlation testing may be overly permissive, with less informative edge weights.

Clauset, Shalizi, and Newman’s method for statistically determining whether a network is ‘scale-free’ showed that the SLTE network is not [25]. On the SLTE network, the result showed alpha = 2.18, which is concordant with power law networks. However, the goodness of fit test using the Kolmogorov-Smirnov statistic returned a p-value of 0.03, indicating that only a small fraction of the simulated scale-free distributions are "close" to the observed degree distribution.

In the rest of the analysis, only the transfer entropy network is used, since it is clear that the correlation-based network is not a super-set of the transfer entropy network, does not agree in the weighting, and is likely overly permissive with regard to active interactions.

### Iteratively solving the influence maximization problem provides a ranking

Using transfer entropy to quantify information flow, if an upstream node transfers information to a downstream node, respecting edge directions, the downstream node is said to be ‘influenced’. The maximization problem is to find a set of nodes, that when treated as information sources, influence the largest proportion of the network.

The Influence Maximization Problem (IMP) was solved over a range of values for *K*, the number of source nodes. The influence score, representing a network cover, increased quickly for small values of *K*, gradually leveling out. Setting *K=45* source nodes (3.9% of the network) produced a maximum network cover of 1064 nodes (92.8%). Beyond *K=45*, the score became saturated (see SI Fig 2).

As the algorithm is stochastic, the range of *K* (from 1 to 45) was run twice and an average count was made on the number of times genes were selected. The ranking produced by each run was highly stable, eliminating the need for a large number of runs. The top ranked gene FKH1, was selected on average 44 times, followed by two genes, GCN5 and RFX1, that were selected on average 43 times. Overall, 49 genes were selected in at least one run.

### Comparing influence to traditional metrics of centrality reveals similarities

To provide a basis for comparison to the ranked influencers, 15 different centrality measures were computed on the SLTE network. The list of centrality metrics can be found in Table 1 along with a brief description. Further description of these metrics can be found in supplementary text. The top 20 influencers and associated metrics are found in Table S2.

**Table 1.**
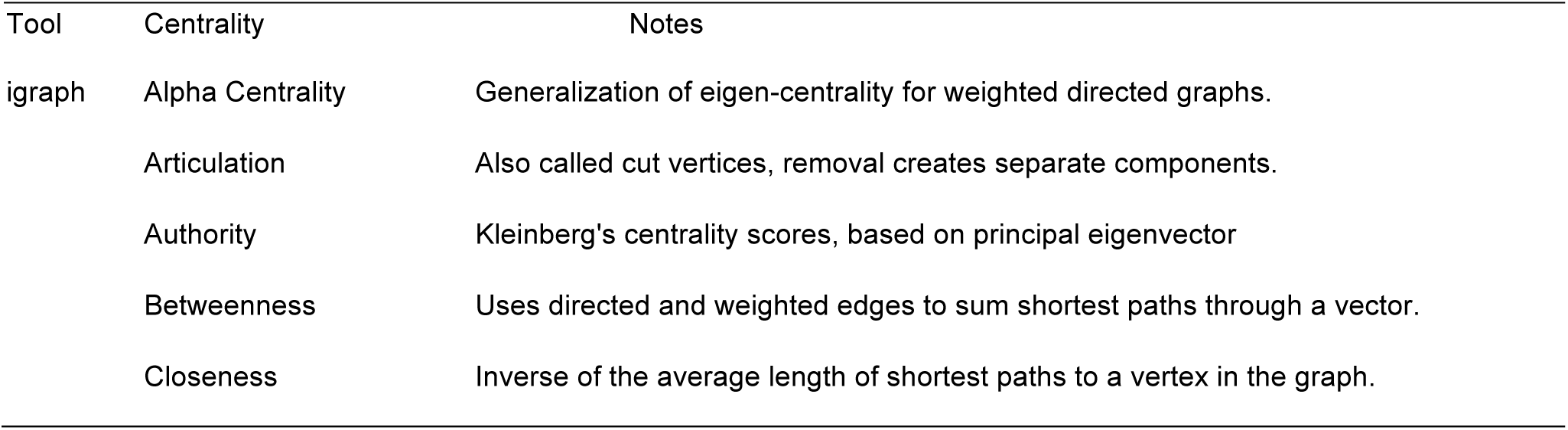
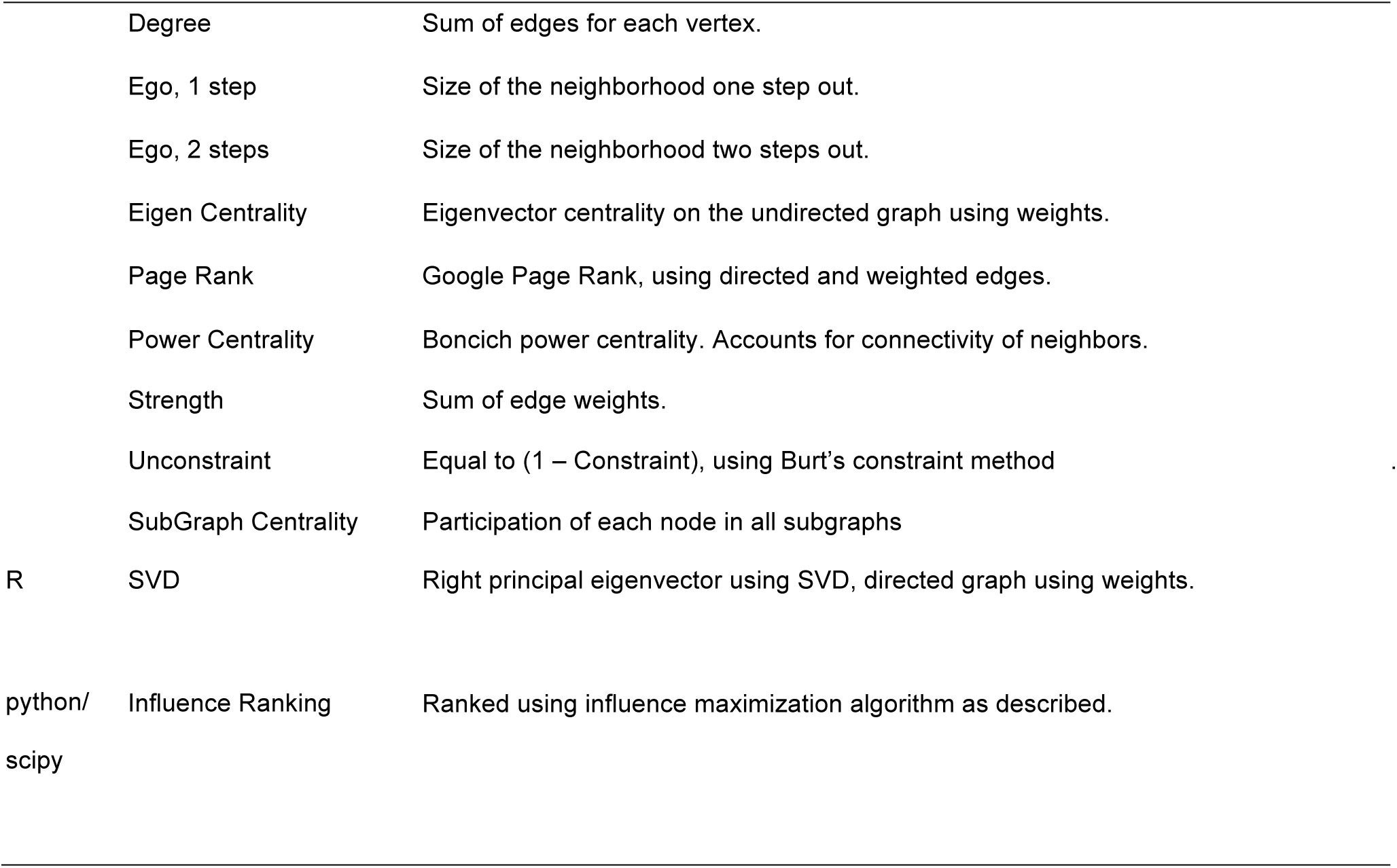
Description of centrality metrics.

| Tool | Centrality | Notes |
| --- | --- | --- |
| igraph | Alpha Centrality | Generalization of eigen-centrality for weighted directed graphs. |
|  | Articulation | Also called cut vertices, removal creates separate components. |
|  | Authority | Kleinberg's centrality scores, based on principal eigenvector |
|  | Betweenness | Uses directed and weighted edges to sum shortest paths through a vector. |
|  | Closeness | Inverse of the average length of shortest paths to a vertex in the graph. |
|  | Degree | Sum of edges for each vertex. |
|  | Ego, 1 step | Size of the neighborhood one step out. |
|  | Ego, 2 steps | Size of the neighborhood two steps out. |
|  | Eigen Centrality | Eigenvector centrality on the undirected graph using weights. |
|  | Page Rank | Google Page Rank, using directed and weighted edges. |
|  | Power Centrality | Boncich power centrality. Accounts for connectivity of neighbors. |
|  | Strength | Sum of edge weights. |
|  | Unconstraint | Equal to (1 – Constraint), using Burt’s constraint method |
|  | SubGraph Centrality | Participation of each node in all subgraphs |
| R | SVD | Right principal eigenvector using SVD, directed graph using weights. |
| python/<br>scipy | Influence Ranking | Ranked using influence maximization algorithm as described. |

Each measure of centrality imparts a ranking over genes in the graph. The top 2.5% of genes was selected from each metric, providing approximately 30 genes for each measure. In metrics with binary values, such as articulation, everything greater than zero was selected. A Jaccard index was computed for each pair of centrality measures (Fig 3). Although some clustering is observed among centrality metrics, especially for node-degree related measures, there remains substantial disagreement in top ranked genes.

**Figure 3.).**
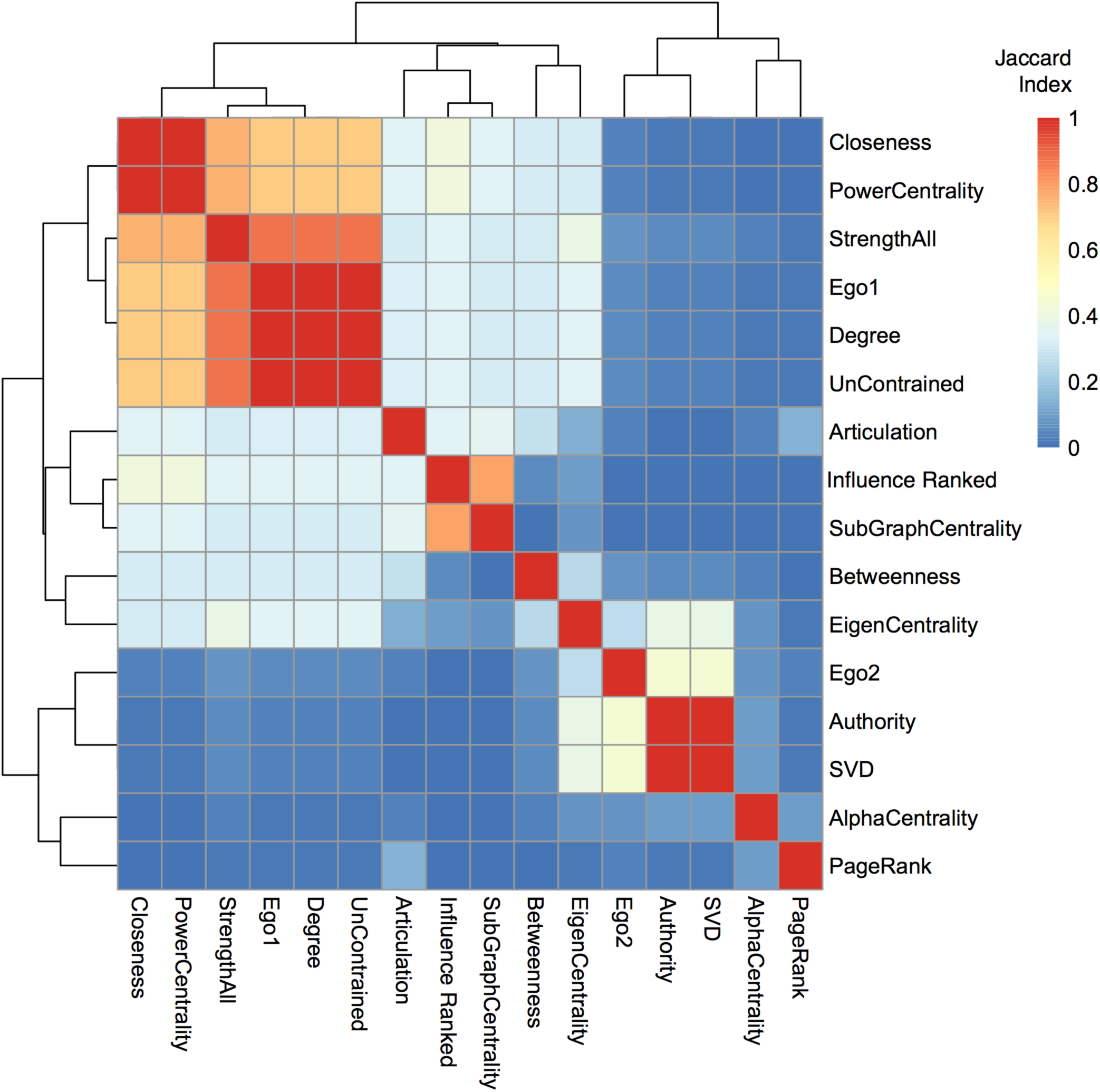
The Jaccard index was used to compare centrality measures. The top 2.5% of ranked genes from 16 different centrality measures were compared using the Jaccard index, which gives values of 1.0 for perfect agreement between sets, and 0 for disjoint sets. There were approximately 30 genes in each set. The dendrogram shows clustering among measures.

The top ranked influential genes are not found among highly ranked genes in eigenvector based centrality measures including authority, alpha centrality, and the SVD derived eigenvector. However, eigenvector measures of centrality contain important genes that are not found in other lists. For example, CLN1 was selected by the SVD derived eigenvector while it was not found on the betweenness list. Overall, no ranked list contained a definitive set of cell cycle related regulators. Across measures, gene set enrichment showed a wide variety of associations with biological processes, illustrating differences in the gene rankings.

**Figure 4.).**
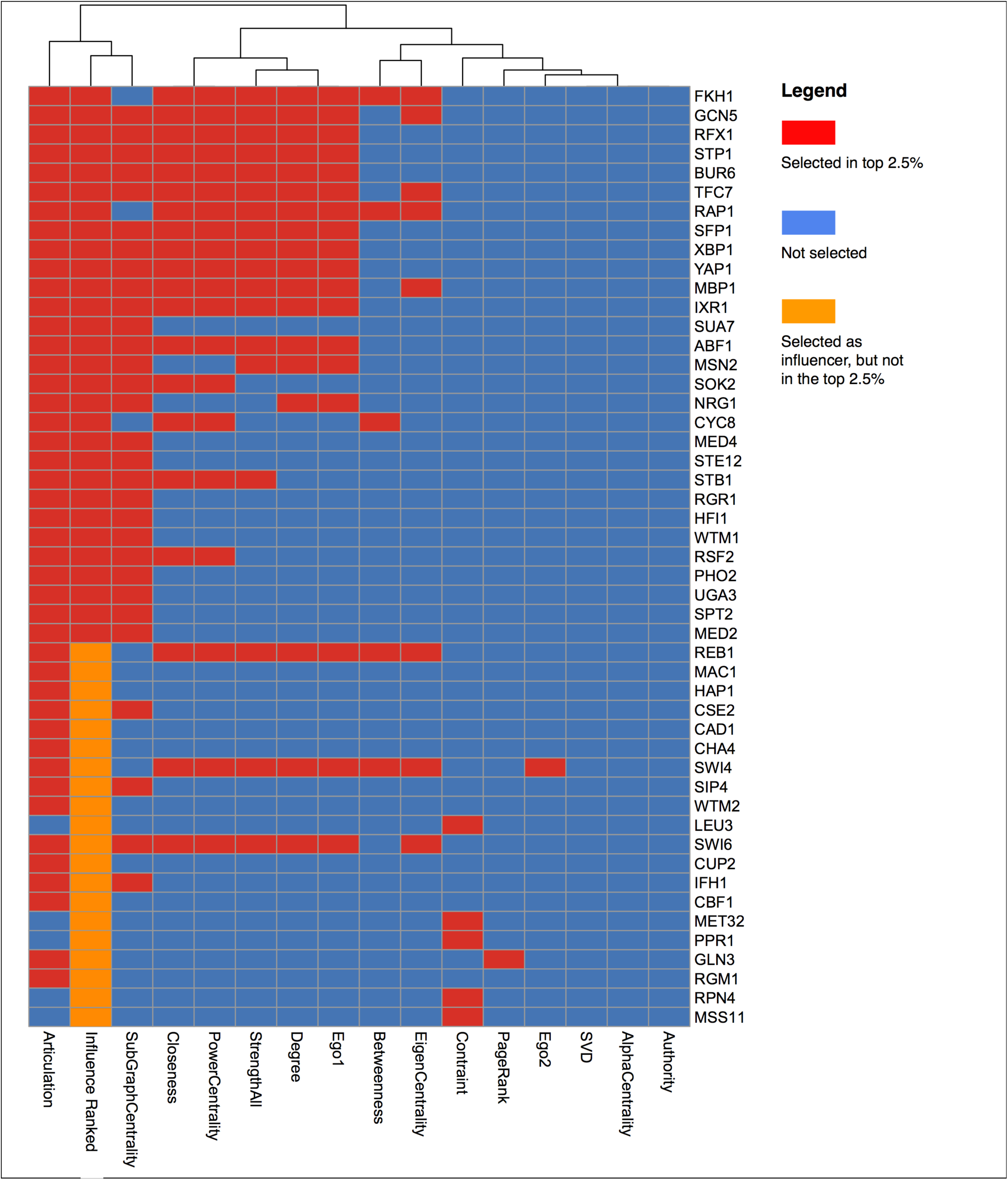
Highly influential genes tend to be selected by other centrality metrics. Genes are sorted by influence ranking in rows (top to bottom), and centrality metrics are found in columns. Genes in orange were influence ranked, but not selected as being in the top 2.5%.

### Influential topology in the regulatory network

We have found that within the regulatory network structure, the influential genes tend to be situated upstream of genes selected by other centrality measures (Fig 5).

**Figure 5.).**
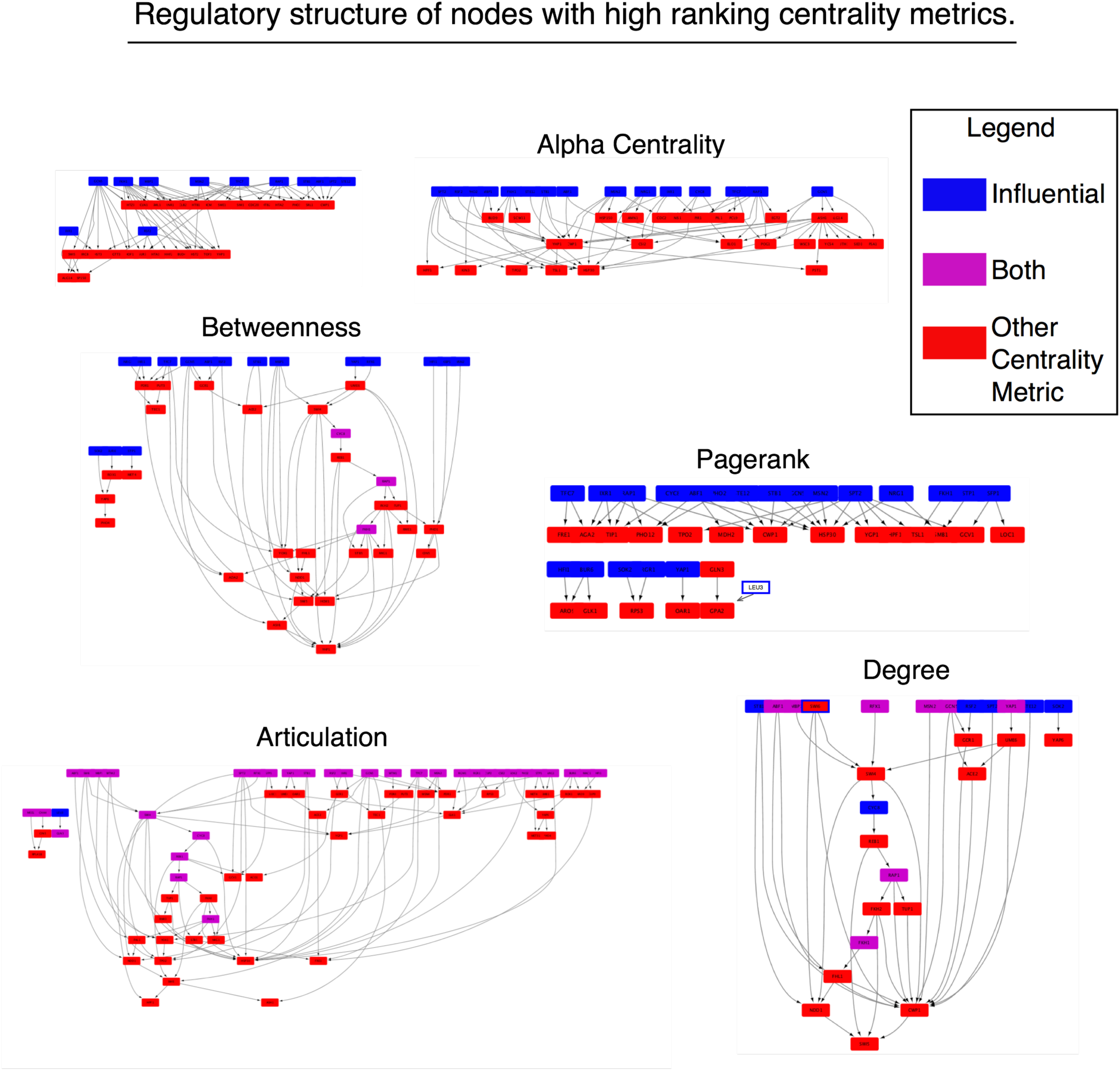
Topology of influential nodes. Highly influential nodes (blue) tend to be upstream of other genes (red) selected by a variety of centrality metrics. Overlapping genes are shown in purple.

For example, the influencer genes act as regulators for genes selected by alpha centrality, while no genes selected by alpha centrality regulate the influencer genes. The same is found for the SVD-derived-eigencentrality and betweenness sets. In some cases, there is a fair amount of overlap in the top-level regulators, such as among the high degree nodes. But, overall, we see the influencers stay as top-level regulators to genes selected by other centrality measures. This can be quantified by computing the fraction of reachable genes, starting at a given measure, excluding overlapping genes (Fig 6). For example, starting at the set of influential genes, 80% of the betweenness selected genes can be reached, while starting at the betweenness genes, only 10% of influencers can be reached. Starting at the influencer genes, 41% of degree central nodes can be reached, while only 10% of influencers can be reached from the degree central nodes. Starting from every centrality measure, the fraction of reachable nodes is less compared to starting from the influential genes. On average, 59% of “central genes” can be reached when starting at the “influential set”, compared to 7% of reachable influential genes, after starting from another measure. The influencer genes are topologically central and connect important genes found by other centrality measures.

**Figure 6.).**
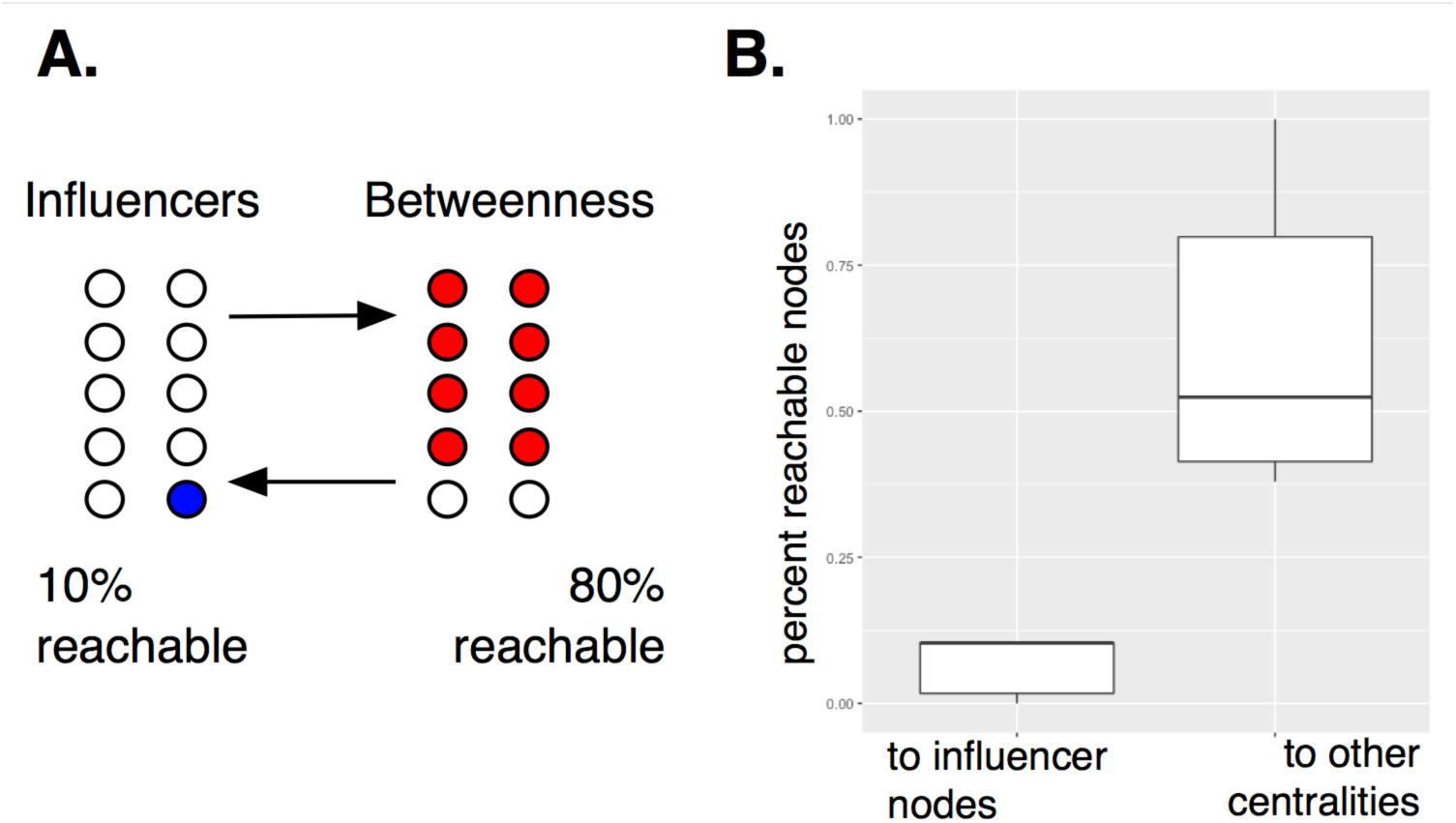
Influence can be quantified by computing node reachability. In (A), an example of node reachability is shown. After starting from a defined set of nodes, *O*, a node, *v*, is considered reachable if there exists a directed edge leading from *O* to *v*. For example, starting at the set of influential nodes, 80% of top ranking nodes using the betweenness measure can be reached, compared to only 10% of influential nodes after starting at the “betweenness nodes”. In (B) node reachability over all centrality measures is aggregated in a boxplot.

In Eser et al., 32 hypothesized cell cycle regulators were named [23]. Comparing the top ranked influential genes, we see again that the influential genes are immediately upstream of the Eser TFs (Fig 7). Although SWI6 was not selected in the top 2.5% of influencers, it is found as a ranked influencer overall. BAS1, on the other hand was not a ranked influencer, and interestingly was not ranked by any other metric of centrality although it has been associated with cell cycle regulation.

**Figure 7.).** Influence ranking of cell cycle related transcription factors. Of the 32 cell cycle related transcription factors given by Eser et al. (red), most are directly downstream of influential genes (blue). Purple shows an overlap between influential and Eser selected genes.

Recently a computational cell cycle model that successfully accounts for 257 of 263 phenotypes [26] was published. In total, 29 genes were extracted from the model where genes making protein complexes were considered separately (SWI6 and SWI4 were used instead of SBF). The full YeastMine network scaffold contained 28 of the 29 genes (CDC55 was not present), and 15 genes were in the SLTE network. Only three genes were ranked as influencers (MBP1, SWI4 and SWI6).

The model is composed of seven modules, each containing between two to eight genes. The MEN module, containing only two genes, was the only module that did not contain any genes from the SLTE network. Among the other six modules, each contained between one to five SLTE genes. While most of the Tyson model genes are not ranked influencers, they are immediately regulated by influential genes. SWE1 is regulated by 8 ranked genes. CDC20 is regulated by 4 ranked genes. CLB5 is regulated by 7 ranked genes. SIC1 is regulated by 5 ranked genes. So in almost all cases, the Tyson model genes are not regulated by a single influencer, but by multiple influencers.

## Discussion

Transfer entropy has been shown to be useful in quantifying information transfer. Here, we show that computing edge weights by summing over time lags, and using a permutation testing framework, leads to biologically salient network structures. Even though the network was constructed by considering all possible regulatory edges, it recovers much of the structure and functional enrichment that one would expect, as demonstrated by the lists of genes returned by commonly used centrality metrics like betweenness and degree.

Since the edges encode dynamics of gene expression by representing information transfer between regulators and targets, flow based methods are particularly relevant. In network flows, some imaginary quantity ‘flows’ from node to node, limited by the capacity of the edges. Here the capacity is represented by the sum of transfer entropies over a small number of time lags. Time lags are important to consider since information transfer is not instantaneous, but instead occurs over a span of time in a biological system. By taking a sum over time lags, all edges are put on the same ‘time frame’, and are thus more comparable.

Edges with the highest weights, implying greatest information transfer, include (FKH1 → HOF1, SLTE=4.79), (SWI4 → HTB1, SLTE=4.75), (FKH2 → IRC8, SLTE=4.47), (SWI4 → SWI1, SLTE 4.34) and (NDD1 → AIM20, SLTE=4.30). The source nodes are well-known, multi-functional, cell cycle related transcription factors. The target nodes have more focused functions. HOF1 regulates portions of the actin cytoskeleton. HTB1 is a histone core protein required for chromatin assembly. IRC8 is a bud tip localized protein, with unknown function. SWI1 is a subunit of the SWI/SNF chromatin-remodeling complex. Finally, AIM20 has unknown function, but over-expression leads to the arrest or delay of the cell cycle.

Some well known cell cycle regulators, such as BAS1, were not selected by any centrality measure. As far as influence is concerned, this can be explained by exploring its neighborhood in the network. In the SLTE network, BAS1 is a source to four other genes, all with no influence ranking and subgraph centrality of 0. Among the four targets is PHD1, also a target from XBP1, which does happen to be a ranked influencer, and happens to have degree 32 and a high subgraph centrality (26.8). So, although BAS1 is probably a cell cycle regulator, there are better sources to choose when targeting the same downstream genes. XBP1 binds cyclin gene promoters and is stress related. Interestingly, it’s important in G1 arrest, which relates to the synchronization method used by Eser et al. in the data generation. XBP1 is also a member of the SWI4 and MBP1 protein family. While BAS1 is involved in biosynthesis pathways for histidine, purine, and pyrimidines, and predicted to be involved in mitotic crossover, XBP1, with regard to cell cycle, is certainly understandable in it’s influence ranking. No one centrality metric was ideally suited towards picking out cell cycle related genes, although the ranked influencers ‘pointed’ to a large proportion of genes selected by other centrality measures.

When we considered the ranking of influential genes, we saw that high-ranking genes were more likely to be ranked high by other centrality metrics. There are several notable exceptions, where REB1, SWI4, and SWI6 were relatively low ranked influencers, but were highly ranked by other metrics. These examples are notable due to their previously known role in the cell cycle and regular inclusion in models. Proteins SWI4 and SWI6 are members of the SBF complex, interacting with the MBF complex (SWI6-MBP1) to regulate late G1 events. REB1, essential in some yeast strains, is shown to act as a link between rDNA metabolism and cell cycle control in response to nutritional stress [27], and is shown to have a significant impact on lifespan [28]. The influence ranking was due to higher ranked influencers being upstream of the three genes in the regulatory network. Therefore, they were only selected as *K*, the set of requested influencers, grew large enough.

Gene set enrichment showed functions related to not only cell cycle, but also chromatin remodeling, stress response, and metabolism (see SI). The top two most influential genes, FKH1 and GCN5 have both been related to life span [29,30,31].

Network control is one goal in the study of dynamic networks [32,33]. Given that influential nodes seem to have a topologically advantageous position, one could speculate that influential genes might be useful selections for network control. Biological events that impact the influential nodes, thereby affecting normal information flow, could have a strong effect on the network, potentially leading to disease states. Discovering the minimum sets of biological entities that hold the greatest influence in the network context could lead to further understanding of how network dynamics is associated with disease.

## Materials and Methods

The methods described here have been implemented in python and are freely available. Run times are kept low by computing the diffusion using sparse matrix linear solvers, and using a multicore-parallel strategy for performing ant optimization. The network weighting, optimization, and diffusion methods are independent, allowing researchers to "mix-and-match" their favorite modules.

### Data sources

Eser et al. [23] generated time series expression data from two replicates of synchronized yeast producing metabolically labeled RNA levels every five minutes over 41 time points. The expression series spans three cell cycles, which progressively dampen in wave amplitude, as yeast synchrony is lost. Using a model for detecting periodicity in gene expression, 479 genes were labeled as statistically periodic. Additionally, 32 transcription factors were predicted to be cell cycle regulators.

YeastMine, the database of genetic regulatory interactions in yeast (May 2015) [22] provided regulatory edges. Using 6,417 yeast genes, 33,809 genetic regulatory edges were collected. Edge weights were computed using a variation of transfer entropy, as described below.

### Computing weights using transfer entropy and time-lagged Pearson correlation

Given two genes are connected by an edge, the edge weight was computed in two ways. First, time-lagged Pearson correlation was used with time lags of 0 to 6 steps (0 to 30 mins.), keeping the maximum. Second, a new variation on time-lagged transfer entropy was used similar to what is described in [34,35], termed sum-lagged transfer entropy (SLTE). TE is computed at each time lag and a sum is taken over the set of time lags. This method avoids making a choice about what time lag to use. Additionally, edge weights in the graph are composed of summed time lags, making them directly comparable. Information transfer, in the genetic regulatory context, is relatively slow and takes place over multiple time steps (each step corresponding to 5 minutes).

Time-lagged Pearson correlation is computed by taking two time series, or numeric vectors ***x*** = {*x*_1_, *x*_2_,…, *x*_*n*_} and ***y*** = {*y*_1_, *y*_2_,…, *y*_*n*_}, and computing the correlation on sub-sequences {*x*_1+*k*_,… *x*_*n*−1_, *x*_*n*_} and {*y*_1_ *y*_2_,… *y*_*n*−*k*_}, where *k* is some integer representing the time lag between variables.

Transfer entropy (TE) is an information theoretic quantity that uses sequence or time series data to measure the magnitude of information transfer between variables [36,37]. Transfer entropy is model-free, directional, and shown to be related to Granger causality [38]. In TE, given two random variables *X* and *Y*, where *X* is directionally connected to *Y* (or *Y* → *X*), we would like to know if prior states of *X* help in the prediction of *Y*, beyond knowing the prior states of *Y*. With some simplifications, transfer entropy is straightforward to compute.

Given two sequences ***x*** and ***y***, we describe transfer entropy as

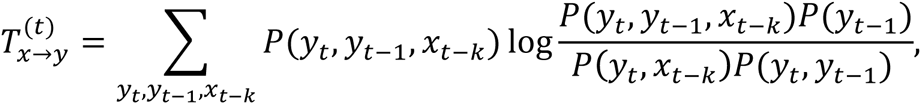

where ***x*_*t*−*k*_** indicates value of the sequence at time step *t*, with time lag *k*.

To perform the computation, ***x*** and ***y*** are normalized and used to fit a Gaussian kernel density estimate (KDE) with the default adaptive bandwidth. Then, by sampling the density estimate, and normalizing the samples, a three-dimensional grid representing the joint probability is generated. The required distributions are marginalized from the joint distribution. Smaller grid sizes provide a finer grained probability distribution, but greatly slow the computation without changing the values substantially. A three dimensional grid of 10^3^ points was found to be a good compromise between computation time and accuracy.

A permutation test is performed to assess statistical significance of the transfer entropy, *T*_***x*→*y***_. The sequence ***x*** is permuted over some number of trials and a count is taken on the number of times the permuted TE is greater than the observed TE, giving an empirical p-value.

### The diffusion model is used to score solutions to the IMP

The information flow model, and most of the nomenclature, is described in [12].

The diffusion models are Markov chains with absorbing states [39]. In the model, vertices are first partitioned into sets S and T. The set S contains sources, generating information, which then flow through the network (nodes in T) until reaching a dead end or absorbing back into S.

The stochastic matrix --defining the probability of moving from one vertex to another-- is defined as

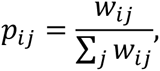

where edge weights *w*_*ij*_ are the weights on outgoing edges. Sets S and T partition the stochastic matrix as

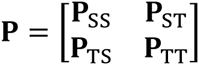

where **P**_SS_ defines the transition probabilities from nodes in S to S, and **P**_ST_ defines transition probabilities from S to T, and so on. Although the matrix is square, it is not symmetric, given the directed edges.

Ultimately, we wish to compute the expected number of visits from a node *v*_*i*_ ∈ S, to a node *v*_*j*_ ∈ T, defined as matrix **H**. By time point *t*, the estimated number of visits from *v*_*i*_ ∈ S to *v*_*j*_ ∈ T is found as

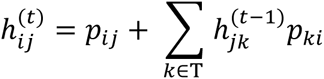

where *p*_*i*_ is the transition probability of *v*_*i*_ ∈ S to *v*_*j*_ ∈ T and 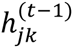 is the expected number of visits that have already taken place at time (*t* − 1), from *v*_*i*_ ∈ S to *v*_*j*_ ∈ T. At time step *t*, information can travel from *v*_*i*_ ∈ S to *v*_*j*_ ∈ T directly, or it would already be at adjacent node *v*_*k*_, and would travel from *v*_*k*_ ∈ T to *v*_*j*_ ∈ T in the next time step. The matrix form of the equation is given as

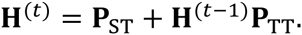

In the long run, at steady state, when **H**^(*t*)^ ∼ **H**^(*t*−1)^, the equation reduces to **H**(**I** − **P**_TT_ = **P**_ST,_ where **I** is the identity matrix. By taking the transpose of both sides, we have (**I** − **P**_TT_)´**H**´ = P´_ST_. This form lets us avoid the matrix inverse in solving for **H**, which can be expensive to compute, and lets us use iterative solvers that can even handle singular matrices. Specifically, the Python SciPy sparse linear algebra library has solvers appropriate for this problem.

To compute a measure of influence on the network, after solving for **H** the expected number of visits on nodes, the influence is summarized as the “influence-score”,

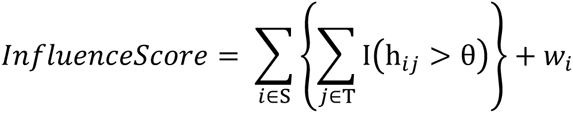

where h_*ij*_ is the number of visitations (using matrix **H**) from node *v*_*i*_ ∈ S to connected nodes *v*_*j*_ ∈ T. Indicator function I(h_*ij*_ > θ) is equal to 1 if the number visitations is greater than a threshold *θ* and *w*_*i*_ is the sum of outgoing weights for node *v*_*i*_ ∈ S. The sum of edge weights is used as a tie-breaker in the case of degenerate solutions, and also makes the case for a solution set that is best supported by data. This influence score is the equivalent to computing the cover on nodes in T. In this work, *θ* = 0.0001 is used.

### Ant optimization is used to search for influential nodes

An implementation of the hypercube min-max ant optimization algorithm was constructed to search for solutions to the Influence Maximization Problem [40,41]. Ant optimization is based on the idea of probabilistically constructing potential solutions to a given problem, in this case a subset selection problem, and reinforcing good solutions with a "pheromone" weight deposited on solution components, ensuring that good solutions become increasingly likely in later iterations.

Since the algorithm is stochastic and results can vary, the optimization is repeated for a defined number of runs. Each convergence takes a number of iterations where ants construct solutions, perform a local search, score the solutions using the influence score, and reinforce the components. As a run progresses, the pheromone values move to either one or zero, indicating whether the component was selected.

At the start of each iteration, ants construct potential solutions, a subset of vertices, by sampling from nodes using probability distribution

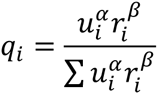

where *q*_*i*_ is the probability for sampling any node *v*_*i*_, with the sum of outgoing edges giving node weight *u*_*i*_ and pheromone weight *r*_*i*_. The alpha and beta parameters are used to give importance to either node weights or pheromones. Solutions are constructed by sampling one node at a time. After each sample, the probabilities are renormalized. Here, *α* and *β* are set to 1.

Local search is performed by stochastic hill climbing, where random, alternative solutions are tried. If a better score is found, the solution is replaced, and carried forward. Local search has a strong effect on the quality of the solutions, and even a small number of hill climbing steps tends reduce the time required for convergence.

Next, using the influence score function, each potential solution is scored, with the best solution kept and compared to solutions found in earlier runs. As part of the Min-Max algorithm, three solutions are kept throughout the run: the iteration-best, the restart-best and the overall-best. The pheromone updates use a weighted average over the three solutions. At the beginning of the run, the pheromone updates are entirely from the iteration-best solution, but gradually, the updates are increasingly influenced by the restart and overall-best solutions, which is done to avoid local minima. The weighted average pheromone would be *r*_*avg*_ = *f*_1_*b*_*i*_ + *f*_2_*b*_*r*_ + *f*_3_*b*_*b*_ where *b*_*i*_ is the iteration best, *b*_*r*_ is the restart best, *b*_*b*_ is the best overall, and fractions *f*_1_ + *f*_2_ + *f*_3_ = 1. The pheromone updates are defined as *r*^(*t*+1)^ = *r*^(*t*)^ + *d*(*r*_*avg*_ − *r*^(*t*)^), where *r*^(*t*)^ is the pheromone weights at time *t*, *d* is the learning rate, and *r*_*avg*_ is the average over the three solutions. Eventually, the pheromone weights become sufficiently close to zero or one, and if all runs are complete, the solution is returned with the influence score.

### Additional ‘off-the-shelf’ analysis

BioFabric, R and R packages igraph, pheatmap and ggplot2 were used for visualization and analysis [42,43,44,49]. Cytoscape 3.3.0 was used for vizualizing graphs [45,46]. Pathway and GO term enrichment was generated using the CPDB from The Max Planck Institute for Molecular Genetics [47]. SciPy was used in the software implementation [48].

## Acknowledgements

Thanks to the Shmulevich Lab and the Institute for Systems Biology.

